# Atomic description of the reciprocal action between supercoils and melting bubbles on linear DNA

**DOI:** 10.1101/2023.06.21.545919

**Authors:** Matthew Burman, Agnes Noy

## Abstract

Although the mechanical response of DNA to physiological torsion and tension is well characterized, the detailed structures are not yet known. By using molecular dynamics simulations on linear DNA with 300 bp, we provide, for the first time, the conformational phase diagram at atomic resolution. Our simulations also reveal the dynamics and diffusion of supercoils. We observe a new state in negative supercoiling, where denaturation bubbles form in AT-rich regions independently of the underlying DNA topology. We thus propose sequence-dependent bubbles could position plectonemes in longer DNA.

---

Inside living beings, DNA is subjected to torsional stress due to the reading of DNA sequence in transcription and replication [1], as well as due to the wrapping around proteins like nucleosomes in eukaryotes [2, 3] and nucleoid-associated proteins in prokaryotes [4, 5]. This stress is relieved by the supercoiling of DNA, which generates intertwined loops (or plectonemes), and changes in molecular twist, promoting the formation of melting bubbles. The two topological parameters, twist (*Tw*) and writhe (*Wr*), are coupled via *Lk* = *Tw* + *Wr, Lk* being the linking number, which is the total number of DNA turns. DNA supercoiling is one of the basic mechanisms that organizes genomes [6] and, due to its importance, is steadily controlled in cells at a superhelical density *σ* = ∆*Lk/Lk*_0_ ≈ −0.07 [2, 3, 7], *Lk*_0_ being the default linking number.

At the same time, DNA is subjected to pulling forces due to routine activities like polymerase work (with ∼25 pN) [8, 9], loop extrusion or liquid phase-separated condensation (the last two with ∼0.6 pN) [10, 11]. DNA nanomechanical properties have been studied *in vitro* by magnetic tweezers, which allows to impose stretching and torsion to a single molecule [12–15]. This experimental setup, together with the development of elastic rod models [16–18], created an initial picture of how the combination of these two forces disturbs DNA. At low stretching forces (F *<* 0.6 pN), the torsion is first absorbed by twist until it reaches a critical superhelical density (*σ*_*c*_) where it is energetically more favorable for the molecule to bend and buckle into a plectoneme, reducing DNA extension and absorbing torsion as writhe. At higher forces values (F *>* 0.9 pN), DNA extension remains relatively constant at negative supercoiling due to the apparition of denaturation bubbles instead of plectonemes [19, 20].

However, little is known about the actual configurations of DNA because they are not directly observable by experiments. Previous coarse-grained simulations revealed that denaturation bubbles are placed at the tip of plectonemes [21–23], instead of being completely independent as was assumed before [14, 15, 17]. Recently, supercoiled DNA was visualized by combining high-resolution AFM and all-atom molecular dynamics (MD) simulations, using DNA minicircles of around 300-400 bp [24]. This study detected the emergence of bubbles independently of tension, induced purely by supercoiling, which suggests that the crosstalk between these and plectonemes might be more complicated than previously thought, opening the question as to which is the impact on linear DNA.

In this letter, we report a comprehensive set of atomically precise MD simulations to investigate the interplay between supercoiled loops and melting bubbles on topologically-constrained DNA under a physiological range of extension and supercoiling (F= 0, 0.3, 0.7 and 0.1 pN and *σ* = 0, ±0.02, ±0.04, ±0.06, ±0.08 and ±1). To this end, we built linear DNA molecules of 300 bp with a randomly generated sequence containing an AT percentage of 49% and different values of twist. The DNA molecules were modeled using Amber18 [25], BSC1 force field [26] and the implicit generalized Born model with GBneck2 corrections [27, 28] at a salt concentration of 0.2 M following our latest protocols [5, 29]. After minimization and thermalization, we performed an equilibration step of 40 ns with restraints on the canonical hydrogen bonds to avoid the premature disruption of the double helix, allowing re-distribution of twist and writhe [5]. Then, the systems were modeled under a series of restraints in order to maintain the torsional stress and to apply a constant pulling force (see Supplementary Material and Figure S1). The production stage was extended to 0.5-2.7 *µ*s depending on their convergence interval, which was monitored by the cumulative end-to-end distance over time (see Figure S2-S12). Overall, we performed 44 different simulations giving a total trajectory length of more than 32 *µ*s. Only the last 400 ns of each simulation were used for subsequent analysis.

Overall, our simulations show good agreement with experimental force-twist curves [12, 14, 15, 30], as they are able to reproduce the reduction of DNA end-to-end distance at low forces, its flattening at F = 1 pN for *σ <* 0, as well as the transition between extended and plectonemic states at the critical tension of 0.7 pN (Figure 1). As in force-extension curves, we observe that *σ*_*c*_ is higher for stronger pulling strains, as applied tension works against shortening DNA [12, 14, 15, 30]. The reproduction of the ‘hat curve’ gives us confidence in the ability of our simulations to give structural insight into how DNA responds to torsion and tension.

**FIG. 1.**
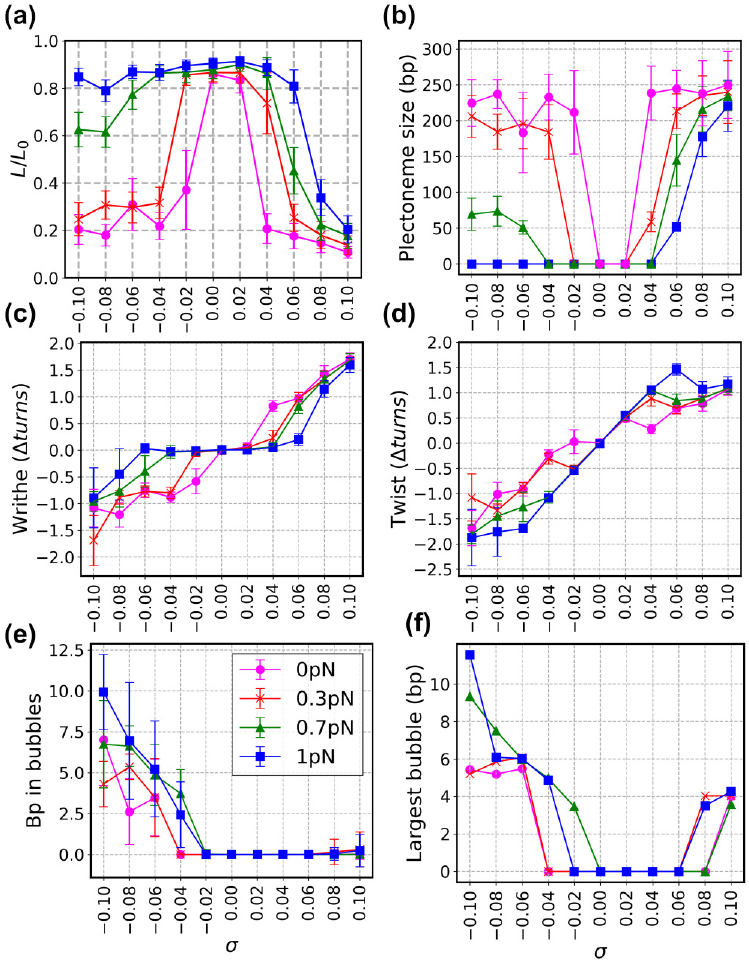
Response of a 300-bp linear DNA molecule to torsion and tension in terms of relative extension *L/L*_0_ (a), plectoneme size (b), writhe (c), twist (d), number of bp in bubbles (d) and size of the largest bubble (e). Values reported here are averages and standard deviations (error bars) over the last 400 ns of each all-atom simulation.

We measured twist and writhe using WrLINE [31] and DNA bending using SerraLINE [24]. Bend angles were calculated using two tangent vectors separated by 16 nucleotides (approximately a DNA helical turn and a half) as a compromise length for capturing the overall bend produced by a melting bubble or by canonical B-DNA. To ensure we were only capturing significant denaturation events, we assumed that a melting bubble was formed by the disruption of at least 3 consecutive bp for more than 1 ns: canonical H-bonds were all broken and angular bp parameters (propeller twist, opening and buckle) were at least 2 standard deviations away from the average obtained on relaxed DNA. The size of supercoils was measured by projecting the trajectories on to the best-fitted plane using SerraLINE [24] and by detecting crossing points formed by two bp placed *<* 3 °A to each other and at least 40 bp apart along the molecular contour.

At low *σ* values, buckled DNA consists of single chiral loops, which resemble the ‘curls’ predicted by the elastic theory that appear before the extrusion of plectonemes [32–34] and that can diffuse over short distances (Figure 2, 3 and Movie S1 in Supplementary Material). When enough torsional stress is imposed (|*σ* | ≤0.08), our modeled DNA presents proper plectonemes (with more than one crossing point) of 8 ± 1 nm of diameter, similar to the ones observed on DNA minicircles [24], suggesting the structures observed there were representatives of what happens in linear DNA. In general, we observe that plectonemes tend to occupy the entire length, so they are pinned at the middle of our DNA sequence (Figure 1b and 2).

**FIG. 2.**
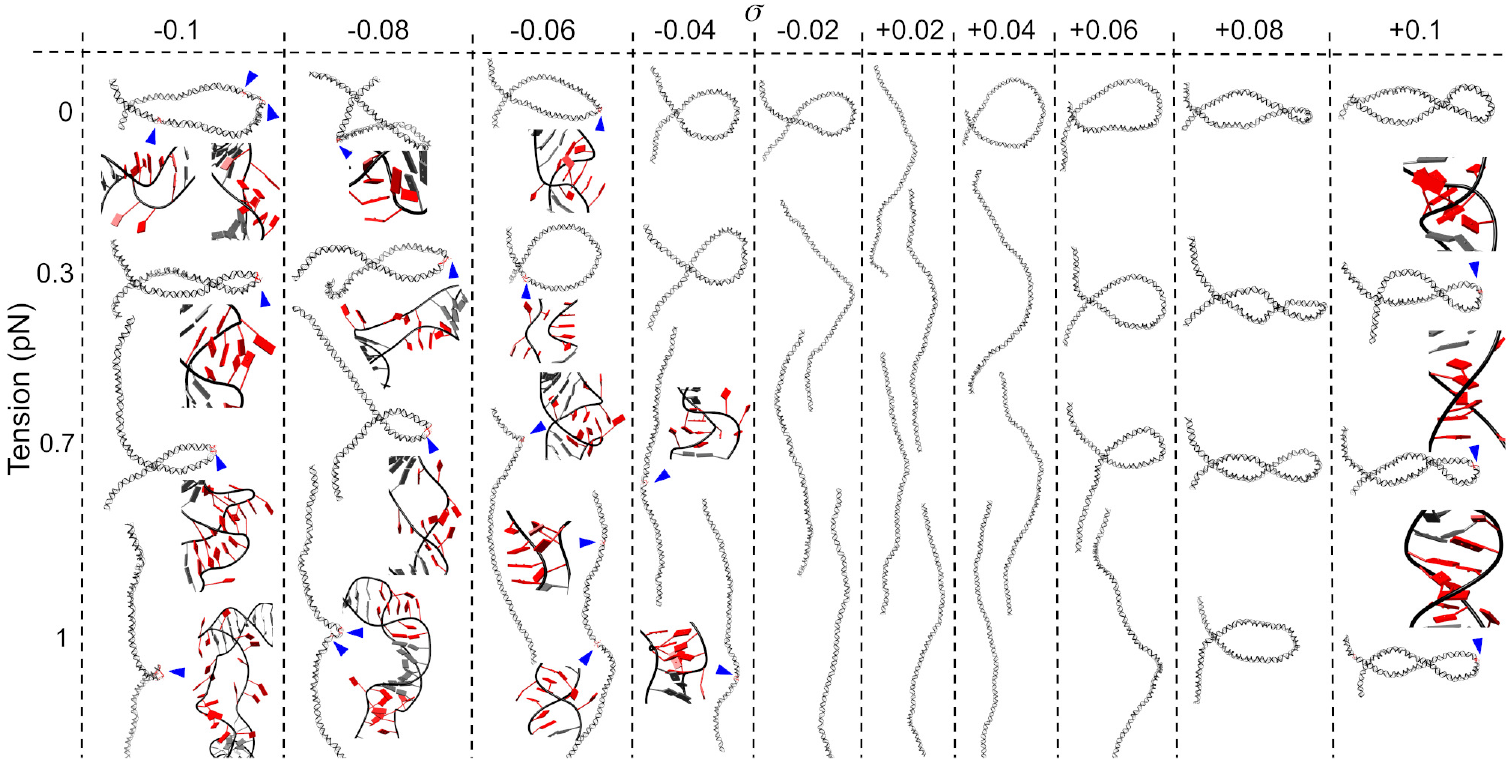
A structural map of DNA under tension and torsion using a representative structure from each of our simulations, where bubbles are highlighted in red and signed by blue arrow heads, with most of them showed in detail by zoomed images.

**FIG. 3.**
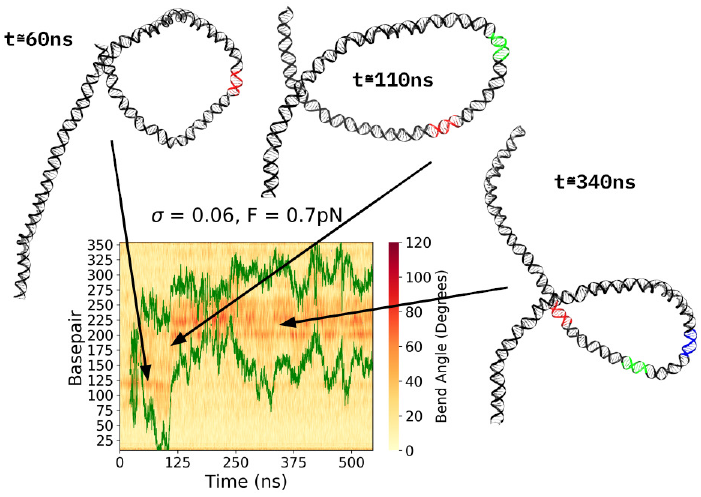
Migration of the supercoiled loop from the simulation at *σ* = +0.06 and 0.7 pN. The kymograph shows loop’s start and end points as green lines and DNA local bends as background heatmap. Representative structures highlight the DNA parts that are sequentially positioned at the tip of the loop: in red at 60 ns, in green at 110 ns and in blue at 340 ns.

For moderate levels of supercoiling (−0.04 ≤ *σ* ≤ +0.06), DNA follows the predictions of simple elastic rod models presenting phase separation between twist-dominated and writhe-dominated states (Figure 1, 2 and Movie S2). In the absence of tension, the formation of negatively supercoiled DNA loops appears at *σ* = −0.02, which is in agreement with previous AFM images on DNA minicircles [24]. At equivalent tensions, *σ*_*c*_ is slightly higher for overwound DNA, compared with unwound DNA, due to the larger capacity of B-DNA for absorbing over-twisting (Figure 1). Buckling is not present in positive supercoiling until *σ* = +0.04 and forms less compacted structures than the ones generated by the same amounts of negative supercoiling, in accordance with previous experiments on circular DNA [35, 36] (see Figure 1a and 2). In addition, the simulations whose end-to-end distances are intermediate also present large fluctuations due to the oscillations between the extended and the buckled phases, as have been previously detected [37, 38] (see Figure 1a, S13 and Movies S1, S3 and S4 in Supplementary Material).

The fact that B-DNA can incorporate relatively high over-twisting, makes the double helix especially resistant to disruption when positively supercoiled. Instead, when DNA is extended in negative supercoiling, its canonical hydrogen bonds and stacking start breaking, forming melting bubbles in order to incorporate the excess of superhelical stress (Figures 1, 2, S4 and Movie S2) [39]. Then, melting bubbles act as flexible torsional spots, assimilating large quantities of twist (Figure 4) [40, 41]. Overall, we observe that, in this low supercoiling regime, DNA follows the predictions of simple elastic rod models where chiral loops and bubbles are excluded from each other.

**FIG. 4.**
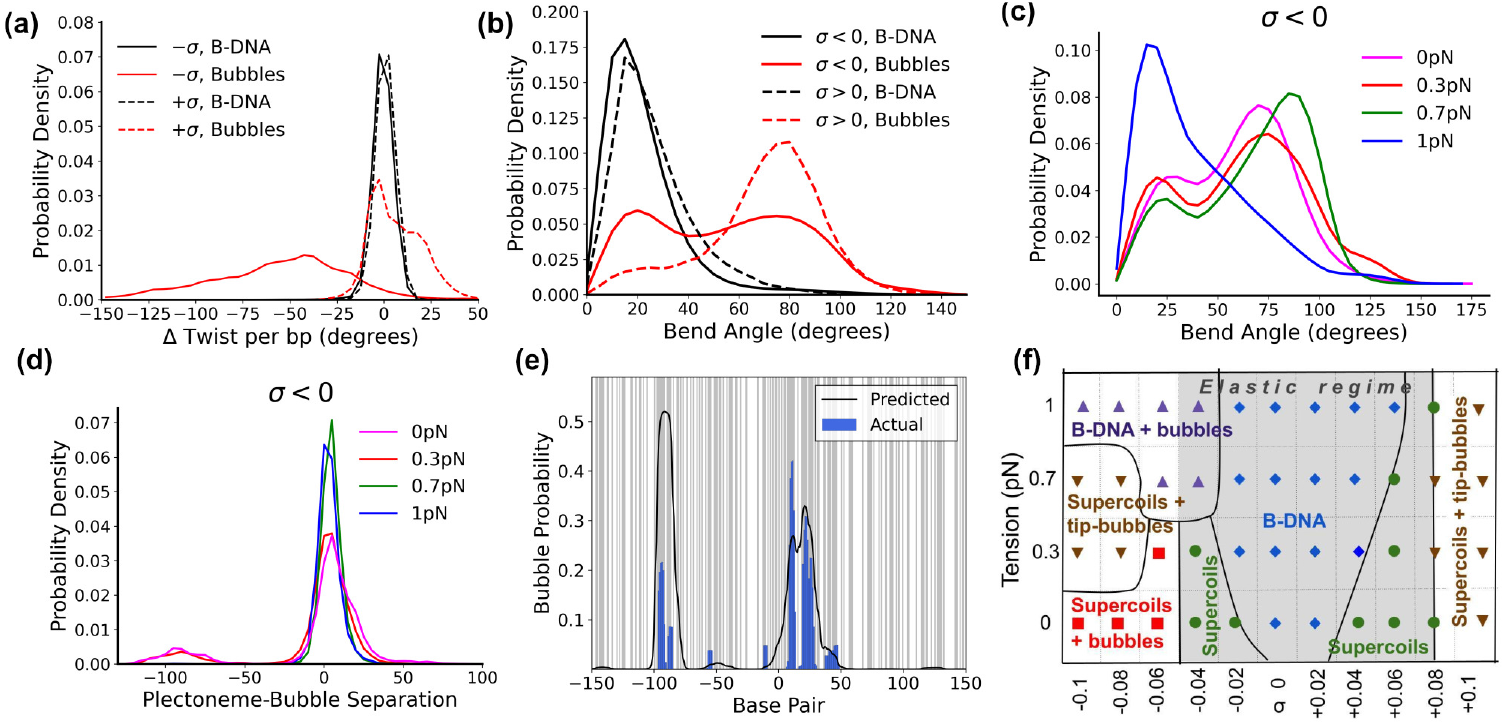
DNA twist (a) and bend angle (b) across all simulations splitted by direction of supercoiling and underlying structure. (c) DNA bend angle of denaturation bubbles formed under negative supercoiling as a function of tension. (d) Distance between the centres of plectonemes and bubbles for negatively supercoiled DNA as a function of tension. (e) Position of denaturation bubbles predicted by the SIDD model (black) and observed in our negative supercoiling simulatioms (blue). A-T bp are shaded in grey. (f) Phase diagram summarising the simulation results showing the various conformational states: extended B-DNA (blue diamonds), B-DNA and bubbles (purple triangles), supercoils (green circles), supercoils and bubbles (red squares) and supercoils with tip-bubbles (inverted brown diamonds). The area where DNA follows the elastic regime is shaded in grey.

On the contrary, DNA under highly negative supercoiling (*σ* ≤ −0.06) presnts most of the melting bubbles at the tip of plectonemes (Figure 4d, S5-S7), because the high curvature there promotes the disruption of the double helix [21, 24]. In addition, the high flexibility of melting bubbles accommodates further sharp bends (Figure 4b), allowing compaction on plectoneme and thus an increase of the end-to-end distance under tension [21]. In agreement with these predictions, we observe plectoneme tip-bubbles to be especially prominent at the critical tension of 0.7 pN (Figure 2 and 4c). Moreover, these ‘tip-bubbles’ are expected to grow when tension is raised, causing shrinking and eventually vanishing of plectonemes [21], as we can see in our simulation at *σ* = −0.1 and 1 pN (Figure S7 and Movie S5). We also observe that the size of the largest bubbles for *σ* = −0.1 increases from ∼5 bp in 0-0.3 pN to around 9 bp in 0.7 pN and 11 bp in 1 pN (Figure 1e).

However, our all-atom simulations reveal a more complex behavior than expected due to the formation of bubbles caused by negative supercoiling and not by tension. At 0 pN, we observe the presence of multiple bubbles in the arms of plectonemes, which can present the same degree of bending as B-DNA (Figure 1, 2, 4c, Movie S6), in agreement with previous observations on circular DNA [24, 42, 43]. Then, for our simulations at *σ* ≤ −0.08, the introduction of pulling tension promotes the concentration of denaturation into a single bubble at the tip of plectonemes, thus converging into the insights given by the oxDNA model [21] (Figure 2 and 4d). At *σ* = −0.06, which is close to the targeted supercoiling level of living organisms [2, 3, 7], we observe an intermediate behaviour between high and low supercoiling regimes: bubbles appear irrespective of tension and tip-bubbles are unessential for the transition into the extended state (Figure 2, S5 and Movie S7). In summary, we show that the DNA structural response for *σ* ≤ −0.06 is highly convoluted as torsional stress can be simultaneously dissipated via plectonemes (or writhe, Figure 1c) and melting bubbles (or twist, Figure 1d, 1e and 4a).

An additional consequence of tension-independent bubbles in negative supercoiling is that their positioning not only depends on the location of plectonemes but also on DNA sequence. As predicted by the SIDD (stress-induced DNA destabilization) program [44], our molecule presents two spots particularly prone to denature due to their high AT content: the first one is close to the center and aligns really well with the tip of plectonemes; the second one is located ∼85 bp off-center and coincides with a small peak of denaturation (see Figure 4e). Interestingly, while this second site is minor in our simulations, it has the highest probability according to the SIDD model. Because this algorithm uses only thermal denaturation energies, it effectively assumes DNA being straight so it does not account for curvature [44]. Hence, our results indicate that, on short DNAs, the location of melting bubbles is balanced between the presence of AT-rich sequences and the location of the tip of the plectoneme, which is forced to be in the middle in our short DNA. For longer DNAs, where several plectonemes can co-exist [32, 45], we anticipate a different scenario: melting bubbles could nucleate the extrusion of plectonemes, making SIDD predictions valid for the location of the latter as well as the former [46].

Our simulations show that highly positively supercoiled DNA (*σ ≥* +0.08) can present denaturation bubbles, even in the absence of tension, in agreement with recent experiments on DNA minicircles [42] (see Figure 1 and 2). In contrast with negative supercoiling, these melting bubbles are smaller, less stable and always placed near the tip of plectonemes (Figures 2, S11, S12 and Movie S8). Because these bubbles are not as efficient as the ones formed in unwound DNA for incorporating excess of twist (Figure 4a), they cannot follow the same mechanism: they cannot grow under tension to eventually substitute the plectonemic loop. Nevertheless, they still bring additional assimilation of superhelical stress in the form of writhe: the accommodation of a strong bend at the tip of plectonemes enables loop tightening, thus allowing further coiling on DNA (Figure 1 and 2, 4b). These bubbles can also present moderate bends in the range of B-DNA when they slightly move from the tip of the plectoneme (Figure 4b, S14 and Movie S8). In general, our simulations show that, in this high positive supercoiling regime, melting bubbles are always nucleated after the formation of plectonemes because they are promoted by high curvature (see Figures S11 and S12). Hence, our study supports the model deduced by Dekker and co-workers, in which the position of plectonemes is controlled by curvature sequence-dependence of B-DNA [47].

## Conclusions

Our simulations successfully connect force-extension curves obtained by magnetic tweezers with microscopy and biochemical data from DNA minicircles, demonstrating that the structural features observed in the latter are also present in linear DNA. Hence, we can provide, for the first time, a phase diagram of the DNA conformation under physiological levels of torsion and tension at atomic resolution (Figure 4f). The elastic regime, where supercoils and bubbles are excluded from each other, is mapped at −0.04 ≤ *σ* ≤ +0.06 and the plectoneme tip bubbles described by coarse-grained simulations are at *σ* ≤ −0.08 and 0.3-0.7 pN. In addition, we uncover two new conformational states: plectoneme tip bubbles for *σ* ≥ +0.08 and melting bubbles nucleated in AT-rich regions for *σ* ≤ −0.06 and minimal tension. At the length of our modeled DNA (300 bp), these sequence-dependent bubbles are not necessarily placed at the tip, as this is constrainted to be in the middle. However, for longer DNA, where plectonemes have room for re-locating, sequence-dependent bubbles could have a key role in nucleating them. We anticipate that these different conformational states and modes will have important implications for the regulation of loops in promoters (with size 100-1000 bp [48]) and of larger topological domains.

## Acknowledgements

This work was supported by the Physical Sciences Research Council (EPSRC) grants EP/N027639/1 (AN) and EP/R513386/1 (MB). The authors acknowledge EP/T022205/1, EP/R029407/1, EP/P020259/1 and the local York facilities for computational resources.

