## Supplementary Material for "Atomic description of the reciprocal action between supercoils and melting bubbles on linear DNA"

Matthew Burman[1] and Agnes Noy[1,*]
[1]*School of Physics, Engineering and Technology, University of York, York, YO10 5DD, UK*

### I. CONTROL OF THE TENSION AND TORSION IN *IN SILICO* LINEAR DNA

DNA was modelled under a series of restraints to control its tension and torsion, thus mimicking force-extension experiments (see Figure S1). These were applied to the cartesian coordinates of a group of selected atoms and to key internal coordinates (also called 'NMR restraints' on the AMBER program [1]). Specifically, one end of the duplex was fixed in place by restraining the coordinates of one atom per each base of the final bp ('fixed end') (atoms I and J in Figure S1a). The other end ('mobile end') was kept under constant tensile force by applying a couple of linear distance restraints between each DNA strand (atoms C and D) and two fixed dummy atoms [2] used as reference points (atoms A and B in Figure S1). The motion of this 'mobile end' was confined to the major molecular axis $Z$ via angular restraints against the two other axis (Figure S1).

In order to prevent the relaxation of supercoiling through the 'untying' of DNA over either end of the molecule, we created an excluded volume response that resembled the presence of a bead typical of the experiments with tweezers. This effect was implemented by enforcing a series of angles ($\psi$ in Figure S1b) to be bigger than 90°, defined per each phosphorus atoms and DNA end. However, one drawback of this approach is that the force pushing a phosphate out of the excluded volume, also pushes the ends of the duplex in the opposite direction, thus counteracting the tensile force. To ensure that these excluded volume restraints were triggered as infrequently as possible, we added 60 GC bp to the mobile end as a buffering molecular stretch. These 'dummy bp' were prevented to bend thanks to a group of dihedral restraints applied to each complementary pair of phosphorus atoms ($\phi$ in Figure S1c). The torsional stress of the DNA molecule was also maintained by ensuring that two representative atoms at each end (at the 61st bp for the 'mobile end' and at the 2nd bp for the 'fixed end') were co-planar with two fixed reference points (see Figure S1a).

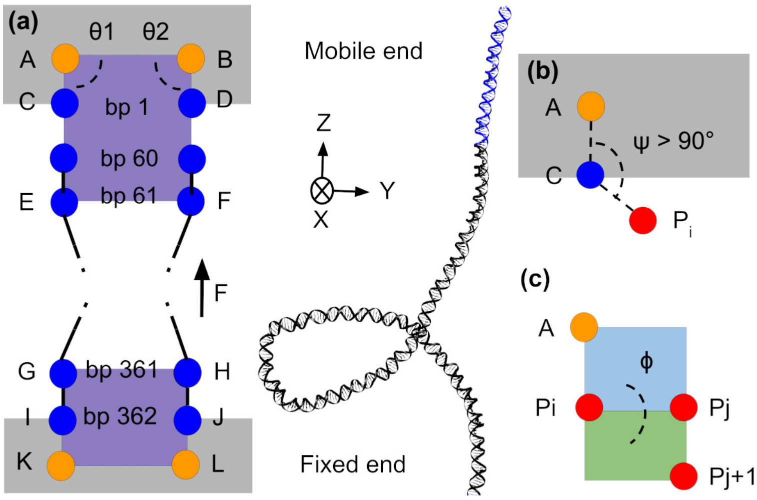


Figure S1. Scheme of the restraints applied in our simulations to control torsion and tension on linear DNA. Dummy atoms, which act as a reference points, are pictured as orange circles. Key O3' or O5' atoms in each ends are pictured as blue circles and phosphorus atoms are pictured as red circles. The motion of atoms C and D is limited to the $Z$ axis, thanks to angular restrains such as $\theta_1$ and $\theta_2$. Grey areas represent excluded volume, in to which the bulk of the strand cannot move. Purple areas represent co-planar planes (ABE -BAF and KLH-LKG) responsible of maintaining the DNA torsionally constrained. The blue and green areas are the two planes that define the dihedral angle $\phi$. The dummy bases in the DNA structure are shown in blue.

### II. SUPPLEMENTARY FIGURES

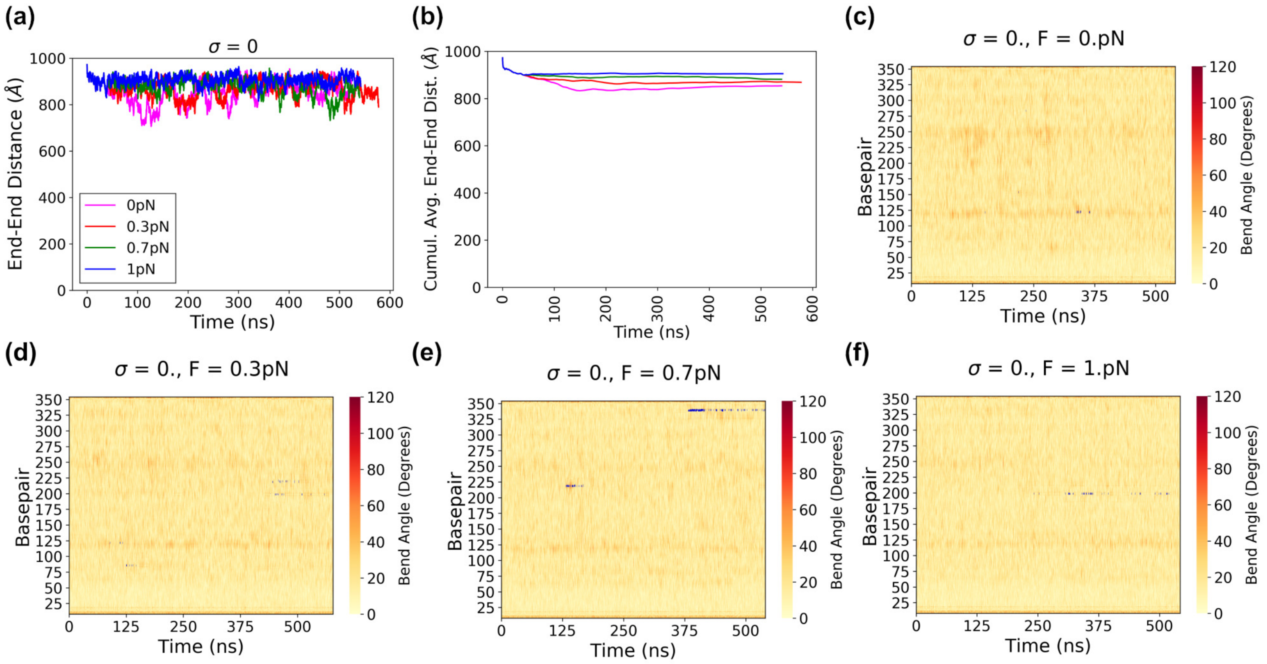


Figure S2. Details of the simulations at $\sigma = 0$. End-to-end distances (a) along with their cumulative averages (b), accompanied by kymographs for each simulation performed at four different tensions: (c) 0, (d) 0.3, (e) 0.7 and (f) 1 pN. Kymographs depict plectoneme boundaries (green lines), denaturation bubbles (in blue) and local bend angles at the fragment length of 16 nucleotides (as a background heatmap). The first 60 bp are the 'dummy bp', which are included as sometimes plectoneme boundaries reach them. Melting bubbles presented here do not account for the condition of being longer than 1 ns.

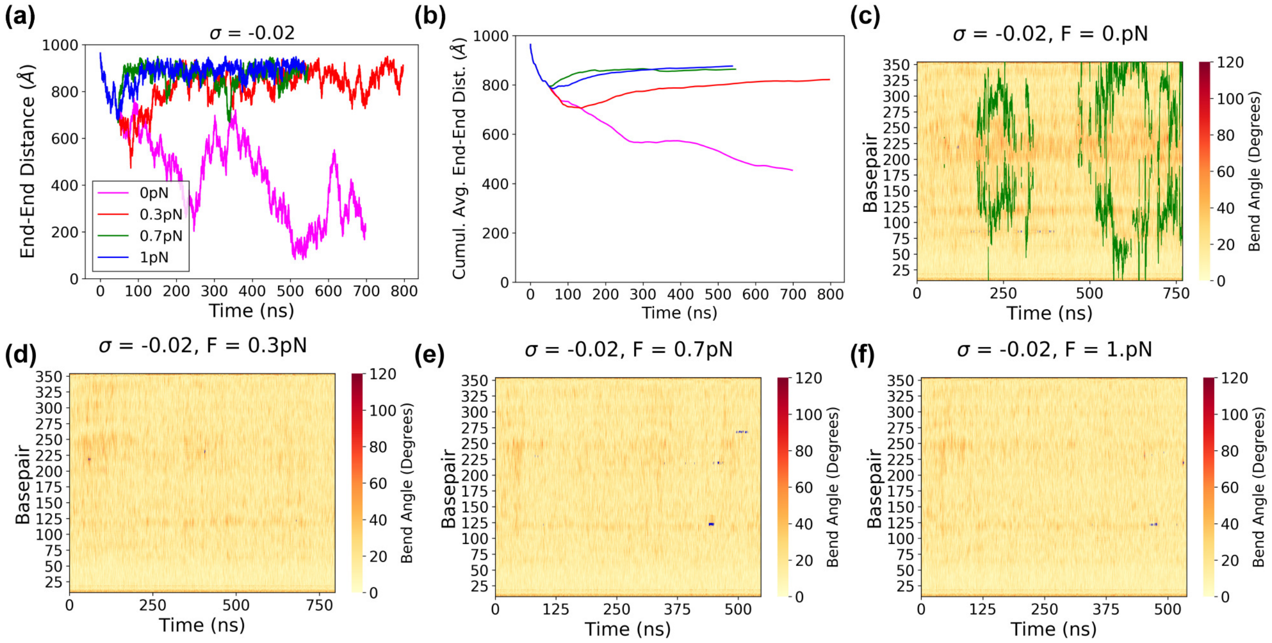


Figure S3. Details of the simulations at $\sigma = -0.02$. Caption as in Figure S2.

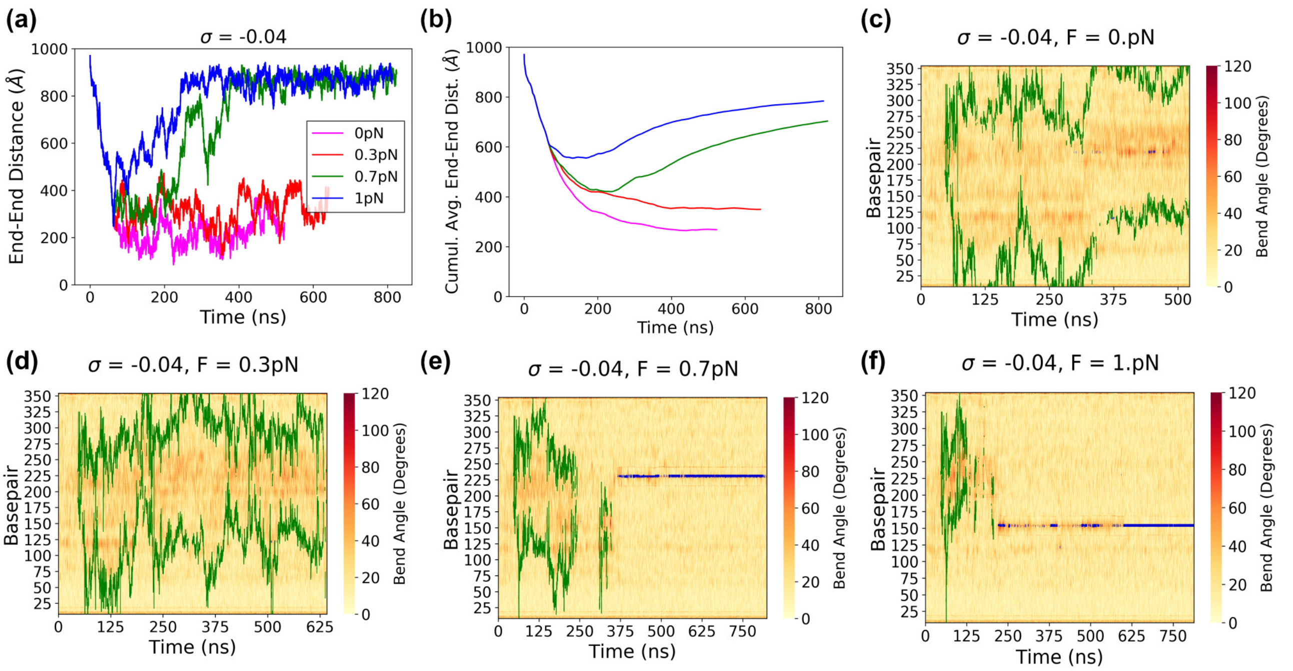


Figure S4. Details of the simulations at $\sigma = -0.04$. Caption as in Figure S2.

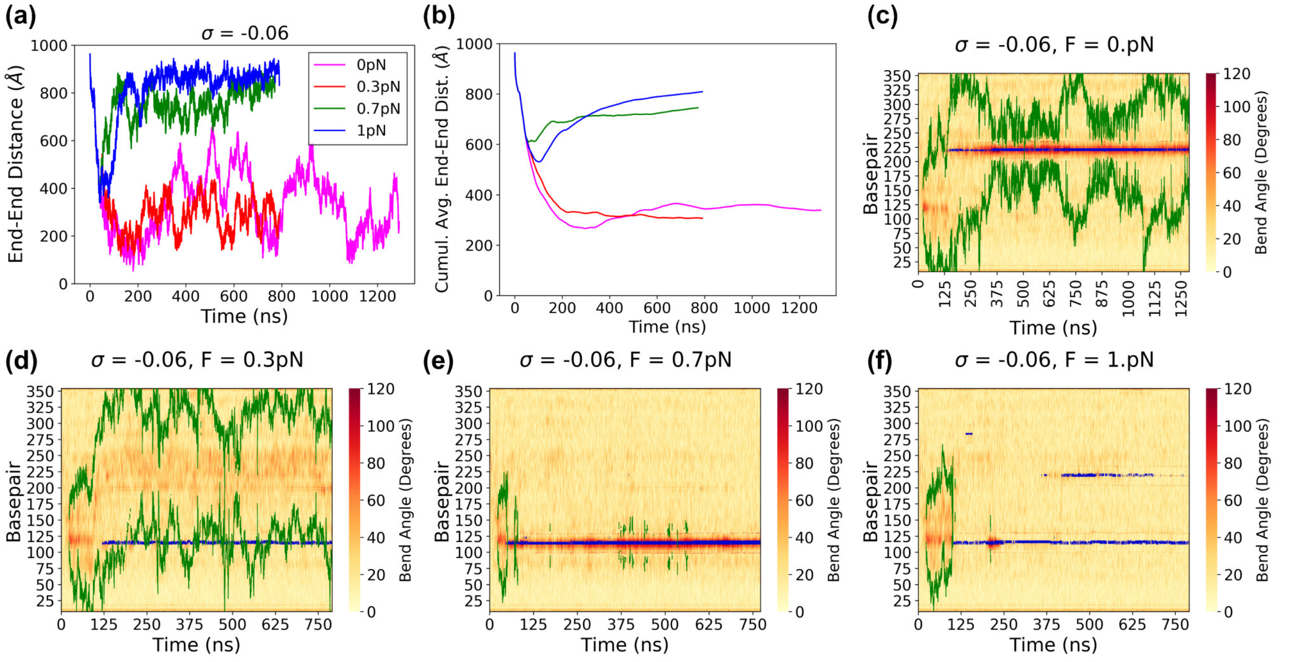


Figure S5. Details of the simulations at $\sigma = -0.06$. Caption as in Figure S2.

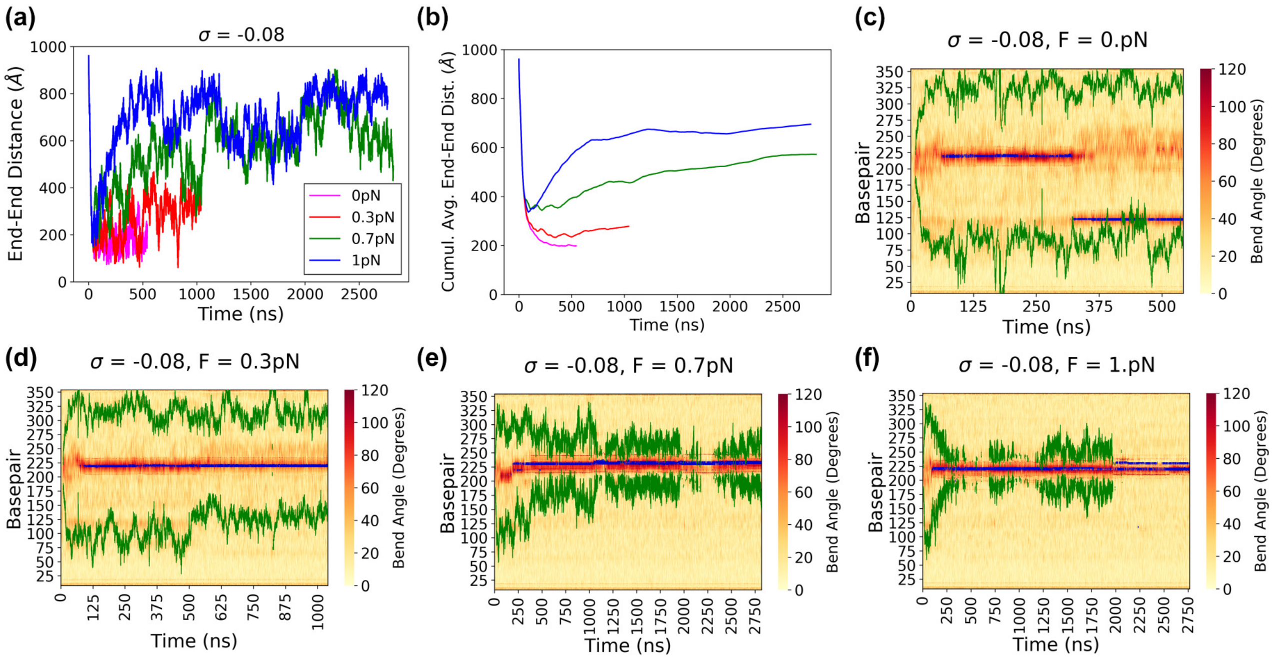


Figure S6. Details of the simulations at $\sigma = -0.08$. Caption as in Figure S2.

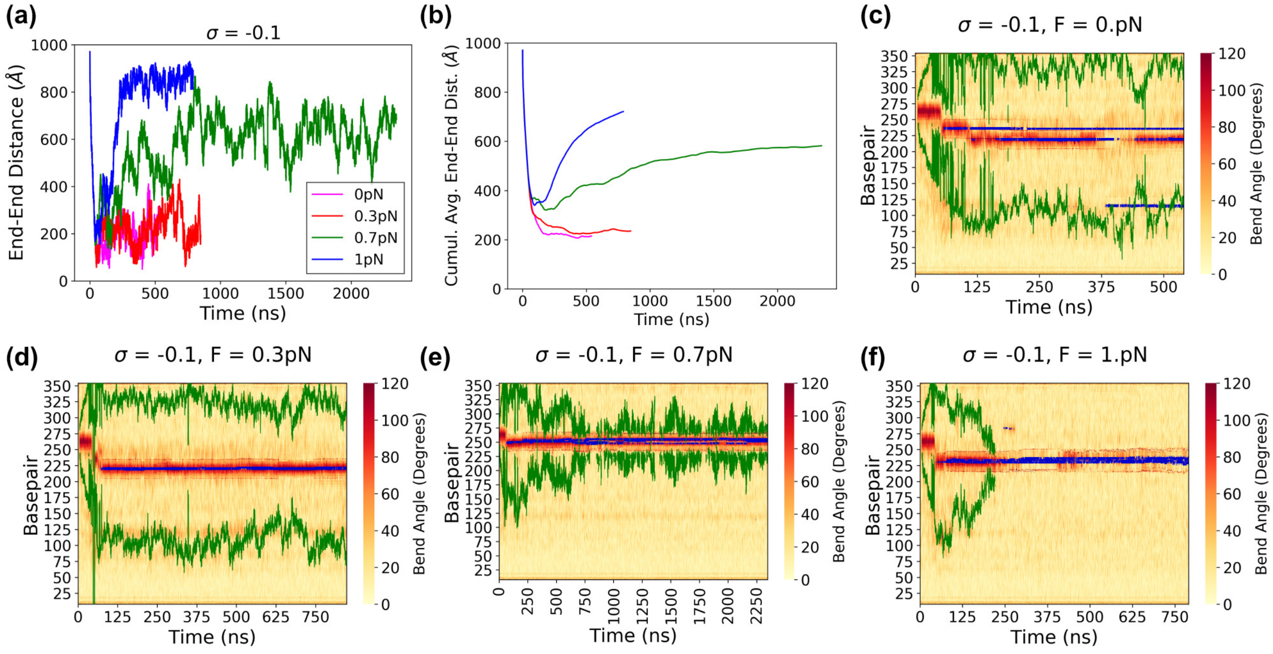


Figure S7. Details of the simulations at $\sigma = -0.1$. Caption as in Figure S2.

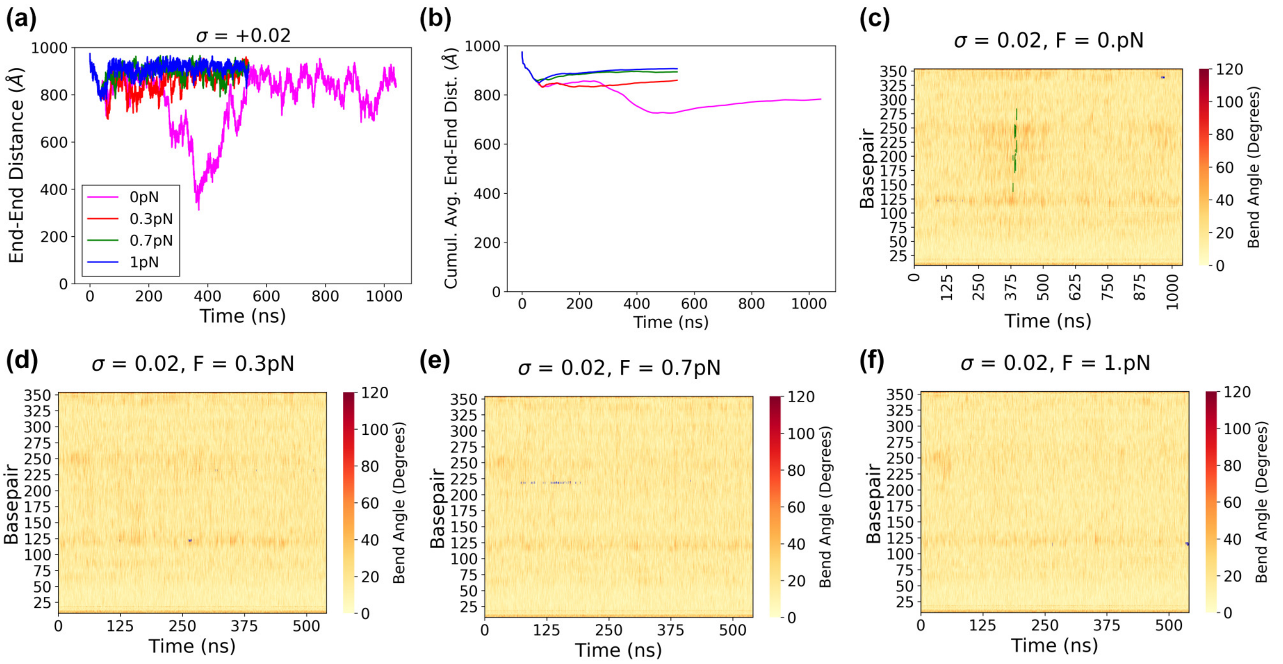


Figure S8. Details of the simulations at $\sigma = +0.02$. Caption as in Figure S2.

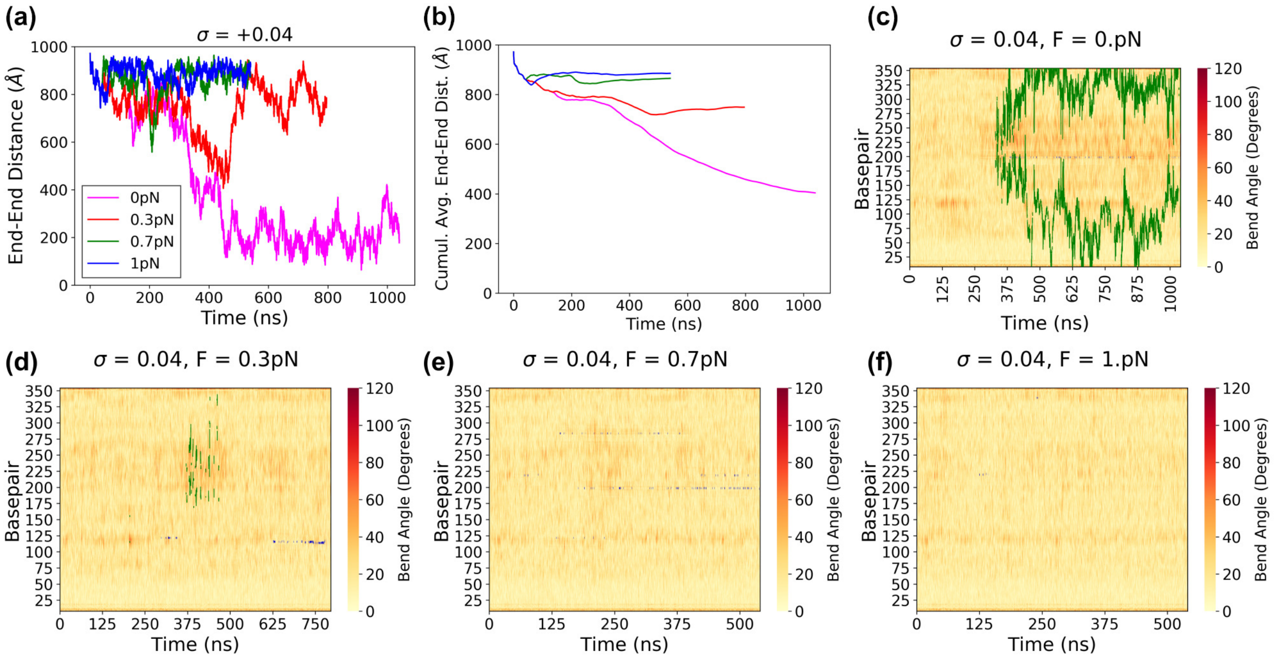


Figure S9. Details of the simulations at $\sigma = +0.04$. Caption as in Figure S2.

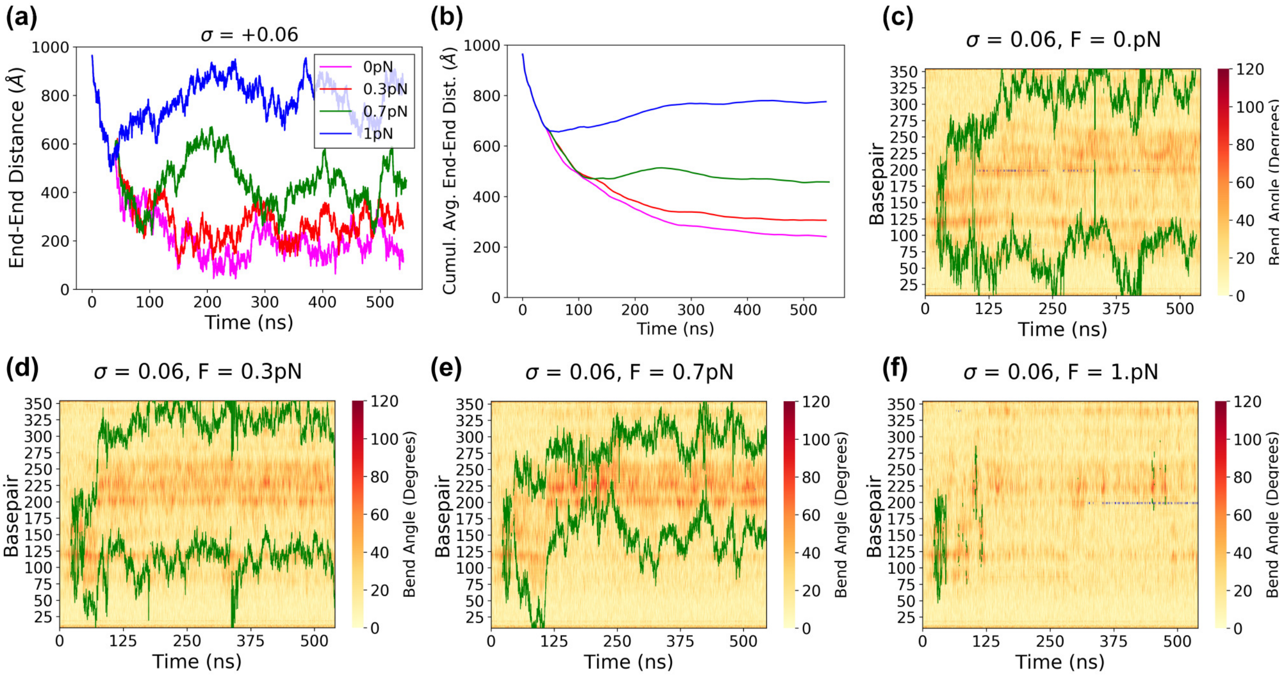


Figure S10. Details of the simulations at $\sigma = +0.06$. Caption as in Figure S2.

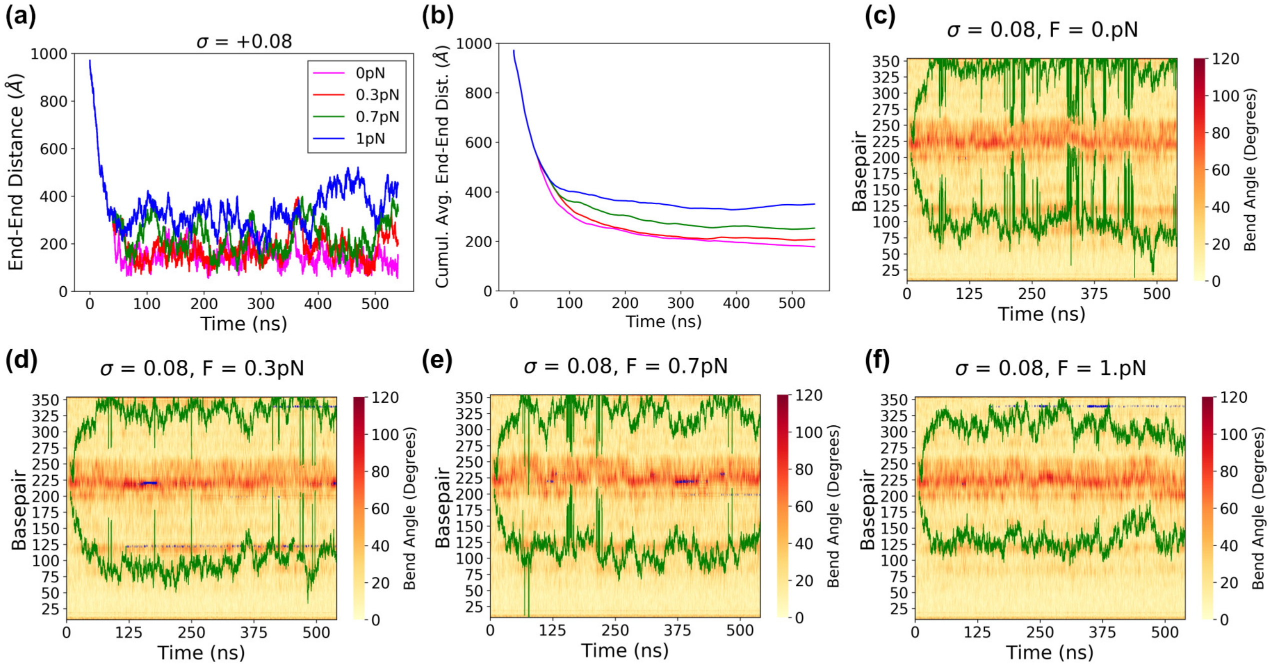


Figure S11. Details of the simulations at $\sigma = +0.08$. Caption as in Figure S2.

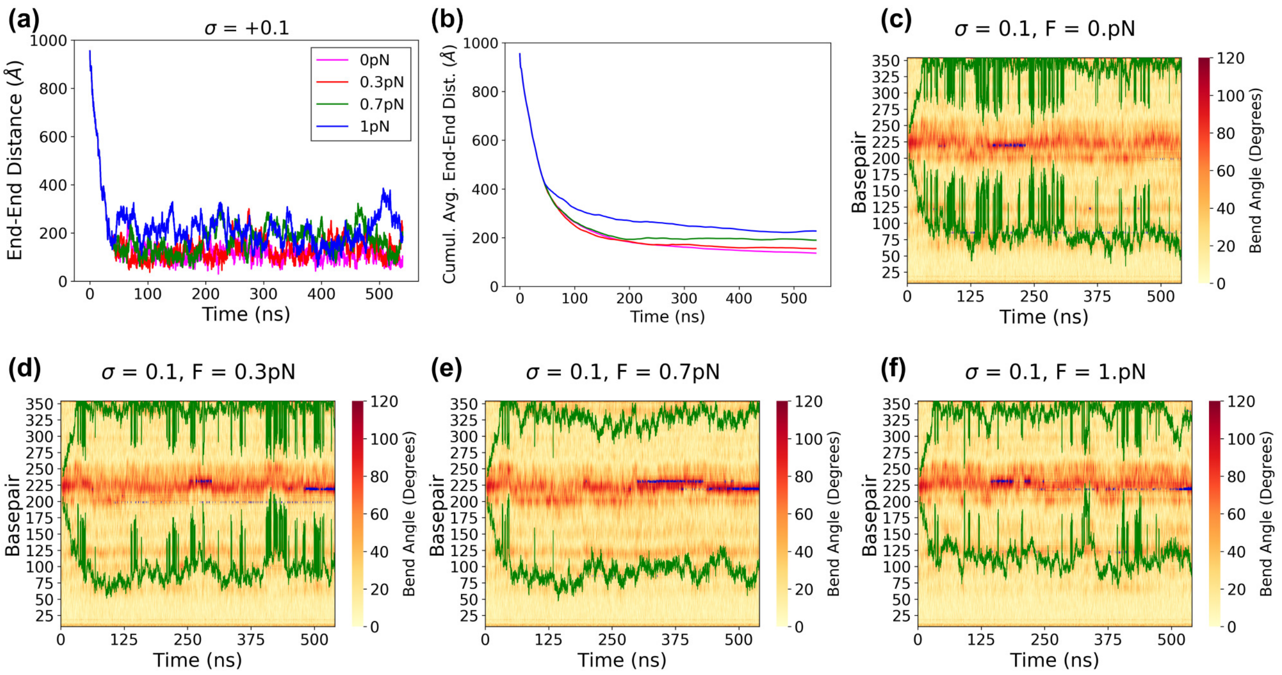


Figure S12. Details of the simulations at $\sigma = +0.1$. Caption as in Figure S2.

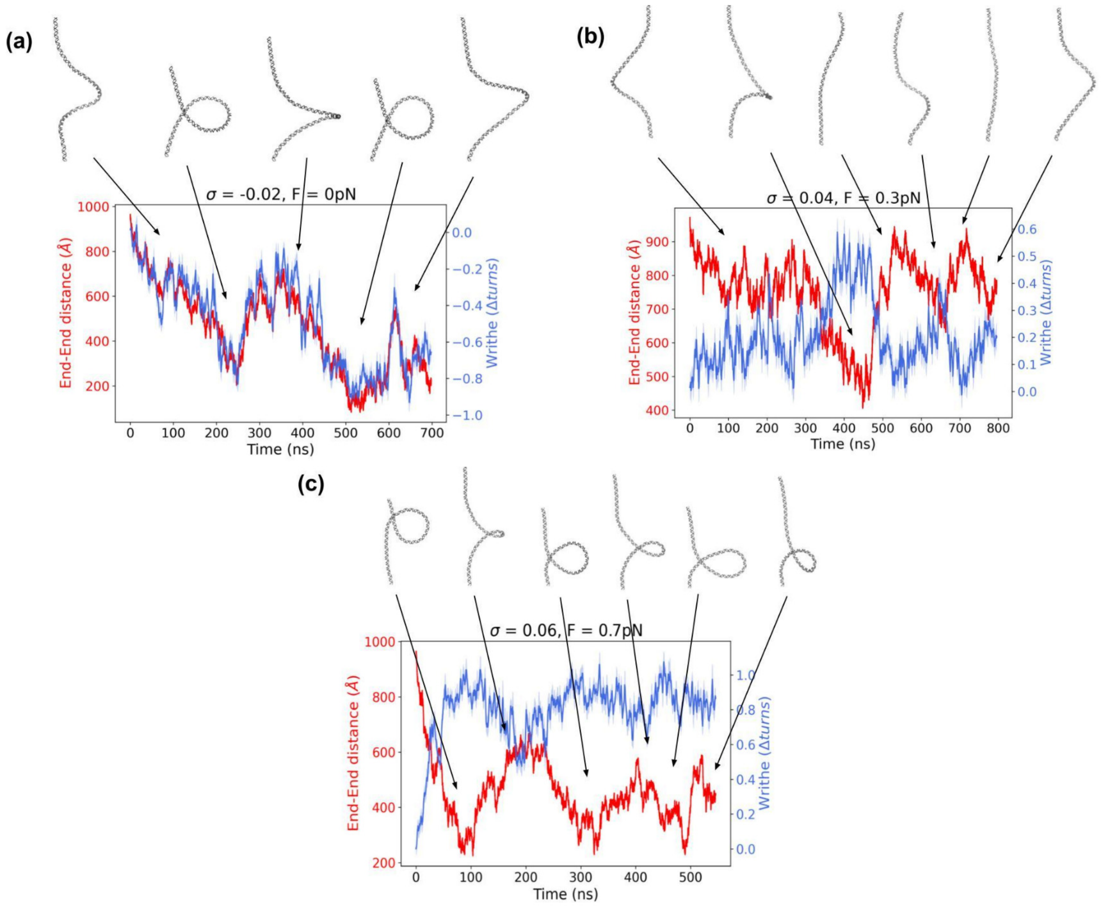


Figure S13. End-to-end and writhe time series together with representative structures showing that big fluctuations in end-to-end distance are due to the reorganization of chiral loops and plectonemes observed in our simulations: (a) $\sigma = -0.02$ at 0 pN, (b) $\sigma = 0.04$ at 0.3 pN and (c) $\sigma = 0.06$ at 0.7 pN

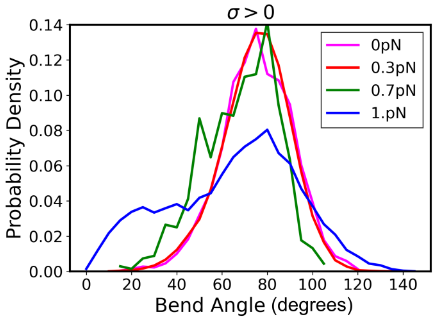


Figure S14. DNA bend angle of denaturation bubbles formed under positive supercoiling as a function of tension.

### III. SUPPLEMENTARY MOVIES

**Movie S1.** Simulation at $\sigma = 0.06$ and 0.7 pN where we can observe migration of a supercoiled loop from one end to the middle and its transient shrinking and expansion, which causes big fluctuations at the end-to-end distance, as predicted by simple elastic theories.

**Movie S2.** Simulation at $\sigma = -0.04$ and 0.7 pN where we can observe a transition from the buckled to the extended state as tension is applied. DNA transforms from a chiral loop where almost all torsional stress is assimilated as writhe, to extended structures where all superhelicity is converted into twist. A melting bubble (in red) is formed due to the exccess of twist in the extended state after plectoneme vanishing, showing separation between denaturation and supercoiled loops at this moderate supercoiling regime which can be described by simple elastic rod models. Denaturation bubbles.

**Movie S3.** Simulation at $\sigma = -0.02$ and 0 pN where we can observe a phase-exchange between the transient formation and vanishing of a supercoiled lopp, which causes big fluctuations at the end-to-end distance, as predicted by simple elastic theories.

**Movie S4.** Simulation at $\sigma = 0.04$ and 0.3 pN where we can observe a phase-exchange between the transient formation and vanishing of writhed structures, which causes big fluctuations at the end-to-end distance, as predicted by simple elastic theories.

**Movie S5.** Simulation at $\sigma = -0.1$ and 1 pN showing the shrinking of a plectoneme and eventually its vanishing at expense of a growing melting bubble (in red).

**Movie S6.** Simulation at $\sigma = -0.1$ and 0 pN showing the formation of denaturation bubbles caused only by supercoiling and not by tension, one of them formed outside the tip of the plectoneme. Only the most long-lived denaturation bubbles are coloured in red.

**Movie S7.** Simulation at $\sigma = -0.06$ and 0.3 pN showing the formation of a denaturation bubble outside the tip of the plectoneme (in red).

**Movie S8.** Simulation at $\sigma = 0.1$ and 1 pN showing the formation of brief denaturation bubbles at the tip of the plectoneme (in red).

[1] D. A. Case *et al.*, "AMBER," (2018), v18.
[2] N. M. Henriksen, A. T. Fenley, and M. K. Gilson, J. Chem. Theory Comput. **11**, 4377 (2015).
